# Caste- and sex-specific DNA methylation in a bumblebee is associated with codon degeneracy

**DOI:** 10.1101/2021.12.17.473163

**Authors:** H. Marshall, M.T. Nicholas, J.S. van Zweden, F. Wäckers, L. Ross, T. Wenseleers, E.B. Mallon

## Abstract

Social insects display extreme phenotypic differences between sexes and castes even though the underlying genome can be almost identical. Epigenetic processes have been proposed as a possible mechanism for mediating these phenotypic differences. Using whole genome bisulfite sequencing of queens, males and reproductive female workers we have characterised the sex- and caste-specific methylome of the bumblebee Bombus terrestris. We have identified a potential role for DNA methylation in histone modification processes which may influence sex and caste phenotypic differences. We also find differentially methylated genes generally show low levels of DNA methylation which may suggest a separate function for lowly methylated genes in mediating transcriptional plasticity. Unlike highly methylated genes which are usually involved in housekeeping functions. We also examined the relationship between the underlying genome and the methylome using whole genome re-sequencing of the same queens and males. We find DNA methylation is enriched at zero-fold degenerate sites. We suggest DNA methylation may be acting as a mutagen at these sites thereby providing substrate for selection via changes in gene transcription mediated by the underlying genotype. However, we did not see any relationship between DNA methylation and rates of positive selection in our samples. In order to fully assess a possible role for DNA methylation in adaptive processes a specifically designed study using natural population data is needed.

## Introduction

In many organisms, the generation of sexual dimorphic traits is mediated via genetically different sex chromosomes. In mammals, this consists of the classic XY systems and in birds, the heterogametic sex is switched, displaying a ZW system (Dean and Mank, 2014). However, the diversity of sex determination systems in insects is considerably more varied. Some species display a X0 system where two X chromosomes are present in females and only one in males (Pal and Vicoso, 2015). Others lack sex chromosomes completely, for example the mealybug, *Planococcus citri*, has a paternal genome elimination system where one whole set of chromosomes is highly condensed in males but not females. A similar system is found in Hymenoptera, haplodiploidy, here males have only a maternally derived haploid genome, generated from an unfertilised egg, whereas females are sexually produced diploids (Normark, 2003). These systems are even more complex in the social Hymenoptera which also display phenotypically different castes of a single sex. The repertoire of molecular mechanisms which allow these species to generate phenotypically different sexes and castes from a single genome remain unknown.

Epigenetic mechanisms have been proposed as a possible method for caste determination in multiple social insect species (reviewed in Sieber *et al*., 2021). The majority of research in this area has focused on DNA methylation, which is the addition of a methyl group to a cytosine nucleotide. Insect DNA methylation, like mammalian DNA methylation, is generally found in a CpG context (referring to a cytosine base immediately followed by a guanine base) (Glastad *et al*., 2014). It is found at lower levels, with <1% - 14% of CpGs being methylated, compared to mammals where around 70% of CpG sites are methylated (Bewick *et al*., 2016; Feng *et al*., 2010). Additionally DNA methylation in insects is generally located in gene bodies and associated with more highly expressed genes, such as housekeeping genes (Provataris *et al*., 2018; Elango *et al*., 2009; Foret *et al*., 2009). The function of DNA methylation in insects is largely unknown and thought to be variable based on the range of overall levels between taxonomic orders (Provataris *et al*., 2018). In Hymenoptera, DNA methylation has been associated with caste differences in various species (Lyko *et al*., 2010; Bonasio *et al*., 2012; Amarasinghe *et al*., 2014; Glastad *et al*., 2016). However, a casual link has yet to be established (Oldroyd and Yagound, 2021). Although no association between the level of sociality of a species and the level of DNA methylation has been found (Weiner *et al*., 2013; Glastad *et al*., 2017).

Many of the studies exploring a role for DNA methylation in the generation of different phenotypes in social insects have focused on single-sex caste differences, although see Glastad *et al*. (2016). However, recently DNA methylation has been strongly implicated in the generation of sex differences in some hemipteran bugs (Bain *et al*., 2021; Mathers *et al*., 2019), with other related species showing minimum sex-specific DNA methylation (Yu *et al*., 2022). This highlights the importance of assessing individual species rather than relying on observations of related species.

Previous work in Hymenoptera has also highlighted a role for the underlying genome in determining DNA methylation profiles. Wang *et al*. (2016) found DNA methylation profiles of alleles in offspring from hybrid crosses of the parasitic wasp *Nasonia* were almost identical to the methylation profile of the parent’s allele. It has also been shown in honeybees that genes which are more variable in terms of DNA methylation show increased genetic variability (Yagound *et al*., 2019) and that DNA methylation marks are heritable (Yagound *et al*., 2020). These studies clearly suggest a link between the epigenome and genome in these species. However, the functional importance of genetically determined DNA methylation remains unknown.

The bumblebee, *Bombus terrestris*, provides an ideal system to assess a potential role for DNA methylation in the generation of sex- and caste-specific phenotypes, in addition to exploring the relationship between the methylome and genome. *B. terrestris* is an important pollinator species both economically and ecologically (Woodard *et al*., 2015). It displays primitive eusociality with colonies consisting of a single female queen, mated by a single male, and female worker daughters. Exhibiting haplodiploidy, male *B. terrestris* are haploid and female queens and workers and diploid. Importantly, genome-wide DNA methylation differences have been found between sterile and reproductive phenotypes of *B. terrestris* workers (Marshall *et al*., 2019), with experimental manipulation of DNA methylation resulting in a change in worker phenotype (Amarasinghe *et al*., 2014). This suggests a functional role for DNA methylation in the generation of phenotypic differences between genetically similar individuals within this species. However, the extent to which DNA methylation differs between sexes and female castes of *B. terrestris* remains unknown. Additionally, previous research has shown colony-specific profiles of DNA methylation in *B. terrestris*, indicating a potential role for genotype-driven DNA methylation profiles in this species (Marshall *et al*., 2019).

In order to determine DNA methylation differences between sexes and castes of *B. terrestris* we have generated whole genome bisulfite sequencing (WGBS) libraries of males, queens and reproductive workers. We have examined the genome-wide methylation profiles of each sex and caste and determined significant differentially methylated genes between castes and sexes. We also carried out whole genome re-sequencing of the same queen and male individuals to determine to what extent the methylation profile of a gene is related to the underlying genotype.

## Methods

### Sample collection

Four colonies of *B. terrestris* were established by Biobest, Leuven. Two colonies were generated from crosses of a *Bombus terrestris audax* queen and a *Bombus terrestris dalmatinus* male and two colonies of were generated from parents of the opposite subspecies, increasing the genetic diversity within our samples. They were then housed at the University of Leuven and kept in 21°C with red light conditions, they were fed *ad libitum* with pollen and a sugar syrup. Callow workers were tagged with numbered disks in order to determine age. Worker reproductive status was confirmed by ovary dissection, ovaries were scored on a 0-4 scale as in Duchateau and Velthuis (1988), entire bodies were then stored at −80°C along with the original queen mothers and male fathers. Three reproductive workers, aged 16-17days, were selected from queenless conditions from each of the four colonies (supplementary 1.0.0).

### DNA extraction and sequencing

Whole genome bisulfite sequencing was generated for the parents and offspring of each colony. DNA was extracted from whole heads of the mother and father of each colony as well as from 12 reproductive workers (three per colony) using the Qiagen DNeasy^®^ Blood & Tissue Kit following the manufacturers protocol. Reproductive workers were chosen to reduce the variation between samples as sterile and reproductive workers show different DNA methylation profiles (Marshall *et al*., 2019). Each sample was treated with RNAse. DNA from the three reproductive worker samples per colony was pooled in equal quantities to produce one representative offspring sample per colony. DNA quantity and quality were determined by Nanodrop and Qubit® fluorometers as well as via gel electrophoresis. Samples were sent to BGI Tech Solution Co., Ltd.(Hong Kong) for library preparation, bisulfite treatment and sequencing. Paired-end libraries (2 × 150bp) were sequenced across two lanes of an Illumina HiSeq 4000 platform with 40% phiX inclusion. A 1% lambda DNA spike was included in all libraries in order to assess bisulfite conversion efficiency, as the lambda genome is known to be unmethylated.

Whole genome re-sequencing of the parents was also carried out. DNA was extracted from half of the thorax of each mother and father per colony following a custom protocol (https://github.com/agdelafilia/wet_lab/blob/master/gDNA_extraction_protocol.md). DNA quan-tity and quality were determined by Nanodrop and Qubit® fluorometers as well as via gel electrophoresis. Samples were sent to Novogene Co., Ltd. for library preparation and sequencing. Paired-end libraries (2 × 150bp) were sequenced on an Illumina HiSeq 4000 platform.

### Generation of N-masked genomes

Differential DNA methylation analyses carried out using samples with different genomic backgrounds, i.e. non-inbred lines, can suffer strongly from reference genome bias, influencing the final differential DNA methylation calls (Wulfridge *et al*., 2019). To account for this we used whole genome re-sequencing data from the parents to call SNPs and create N-masked genomes per replicate colony. Whole genome re-sequencing data of the parents were checked using fastqc v.0.11.5 (Andrews, 2010) and aligned to the reference genome (Bter_1.0, Refseq accession no. GCF_000214255.1, (Sadd *et al*., 2015)) using bowtie2 v.2.2.6 (Langmead and Salzberg, 2013) in *–sensitive* mode (supplementary 1.0.1). Aligned reads were deduplicated using GATK v.3.6 (McKenna *et al*., 2010). SNPs were called using Freebayes v.0.9.21.7 (Garrison and Marth, 2012) which accounts for ploidy differences between males and females. SNPs were then filtered using VCFtools v.0.1.16 (Danecek *et al*., 2011) with the following options: *–max-alleles 2 –minQ 20 –min-meanDP 10 –recode –recode-INFO-all*. A custom script was then used to filter SNPs to keep only homozygous alternative SNPs which are unique to either the mother or father of each colony. We also removed C-T and T-C SNPs as these are indistinguishable from bisulfite converted bases in WGBS. This left a mean of 365,372 SNPs per colony (supplementary 1.0.2). The parental SNPs were then used to create an N-masked genome for each colony (four total) using the BEDtools v.2.28.0 *maskfasta* command (Quinlan and Hall, 2010).

### Differential DNA methylation between castes and sexes

Whole genome bisulfite sequencing (WGBS) data of the parents and pooled worker offspring were checked using fastqc v.0.11.5 (Andrews, 2010) and poor quality bases were trimmed using cutadapt v.1.11 (Martin, 2011). Libraries were then aligned to the colony-specific N-masked genomes created above using Bismark v.0.16.1 (Krueger and Andrews, 2011) and bowtie2 v.2.2.6 (Langmead and Salzberg, 2013) with standard parameters (supplementary 1.0.1). Bismark was also used to extract methylation calls, carry out deduplication and destrand CpG positions. Coverage outliers (above the 99.9th percentile) were removed along with bases covered by less than 10 reads. The methylation status of each CpG was then determined via a binomial model, where the success probability is the non-conversion rate determined from the lambda spike. CpGs were classed as methylated when the false-discovery rate corrected p-value < 0.05. CpG sites were then filtered to remove any site that did not return as methylated in at least one sample.

Differential methylation was assessed at the CpG level in pair-wise comparisons (queen-male, queen-worker, male-worker) using the R package methylKit v.1.16.1 (Akalin *et al*., 2012). A logistic regression model was applied to each comparison with Benjamini-Hochberg correction for multiple testing (Benjamini and Hochberg, 1995). For a CpG to be differentially methylated a minimum difference of at least 10% methylation and a q-value of <0.01 were required. Genes were determined as differentially methylated genes if they contained an exon with at least two differentially methylated CpGs and an overall weighted methylation (Schultz *et al*., 2012) difference across the exon of >15%. Two CpGs were chosen based on Xu *et al*. (2021), they find the methylation of two CpGs is enough to promote gene transcription in *Bombyx mori* via the recruitment of histone modifications.

### Relationship between the genome and methylome

Using the SNPs called above we created alternate reference genomes for the males and queens using the *FastaAlternateReferenceMaker* command from GATK v4.1.9.0 (McKenna *et al*., 2010). We then identified the codon degeneracy of every CDS sequence in each queen and male genome following Mongue *et al*. (2019). Briefly, all codons were labelled per CDS and each base within each codon was labelled between 0-4 depending on how many nucleotide substitutions would be synonymous. We then determined the proportion of zero-fold and four-fold degenerate sites which were classed as methylated (determined via the binomial test above). We also checked the methylation level of a gene against the pN/pS (non-synonymous polymorphisms to synonymous polymorphisms) ratio of that gene (calculated using all genomes in a custom R script) to see if DNA methylation is enriched/depleted in genes potentially under selection. Significant relationships were determined using linear models in R, with interaction effects tested via two-way ANOVAs and post-hoc testing carried out using the *glht* function in the multcomp v.1.4 package (Hothorn *et al*., 2008).

### Gene ontology enrichment

Gene ontology (GO) terms for *B. terrestris* were taken from a custom database made in Bebane *et al*. (2019). GO enrichment analysis was carried out using the hypergeometric test with Benjamini-Hochberg (Benjamini and Hochberg, 1995) multiple-testing correction, q <0.05, implemented from the R package GOStats v2.56.0 (Falcon and Gentleman, 2007). GO terms from differentially methylated genes between sexes and castes and GO terms associated with highly methylated genes were tested against a GO term database made from the GO terms associated with all methylated genes. Genes were determined as methylated if they had a mean weighted methylation level greater than the bisulfite conversion error rate of 0.05 in either queens, males or workers. GO terms associated with hypermethylated genes in any given sex/caste were tested for enrichment against GO terms from the given comparisons differentially methylated genes. REVIGO (Supek *et al*., 2011) was used to generate GO descriptions from the GO ids.

## Results

### Genome-wide sex- and caste-specific DNA methylation

Here we examine the first genome-wide DNA methylation profiles of *B. terrestris* males and queens and compare these profiles to reproductive workers. We find low genome-wide DNA methylation levels in both sexes and castes, similar to those previously reported in workers by Bebane *et al*. (2019) and Marshall *et al*. (2019). On average 0.23% ± 0.05% of CpGs are methylated across all samples, with little overall variation in genome-wide levels between sexes and castes (supplementary 1.0.1). Reproductive workers, queens and males do, however, show different CpG methylation profiles, with males showing more variation between samples and also clustering away from the two female castes (Fig.1a).

**Figure 1:**
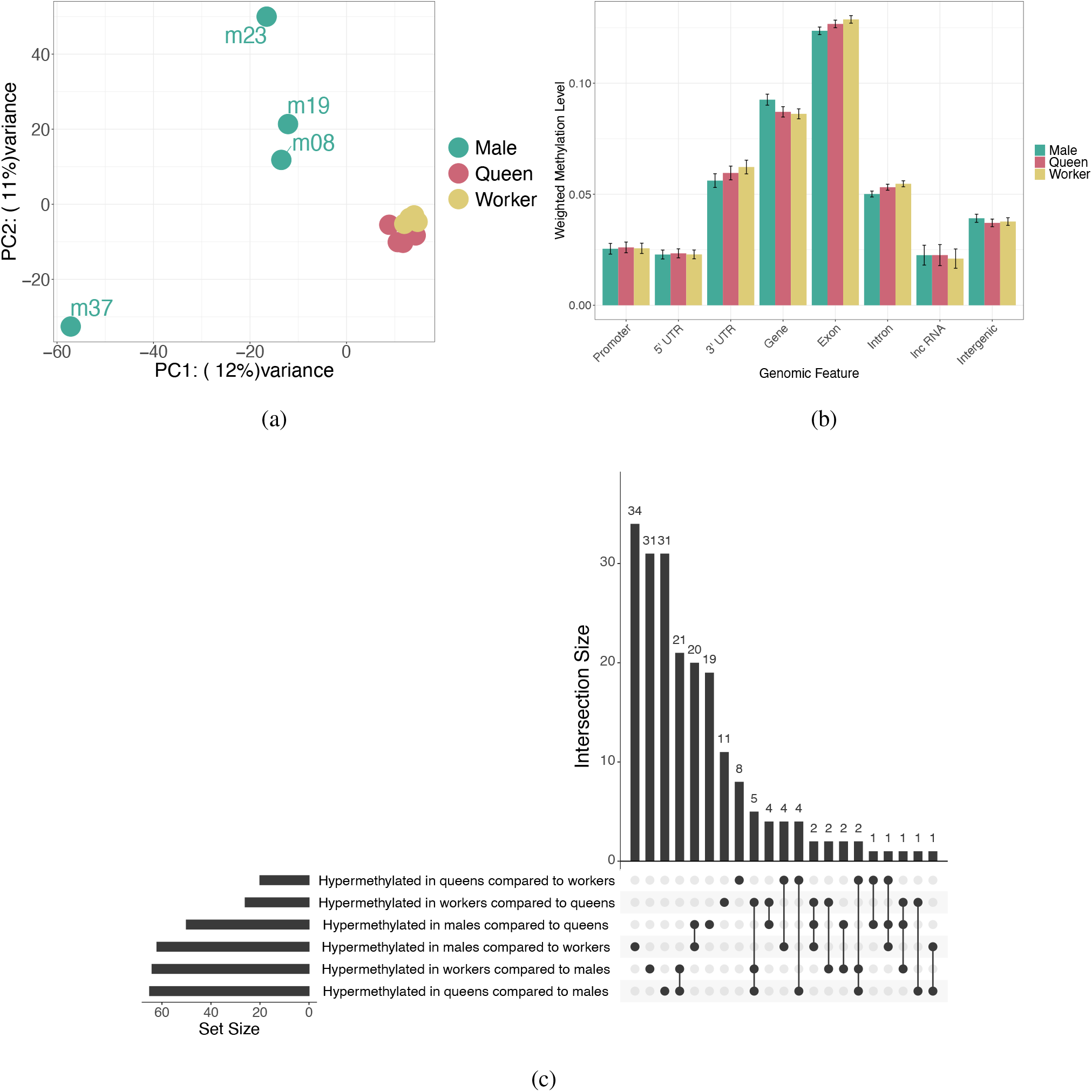
(a) PCA plot based on the methylation level per CpG for all CpGs which had greater than 10X in all samples and were classed as methylated in at least one sample (n = 5,304). (b) Bar plot of the mean methylation level of each genomic feature for sexes and castes. Error bars represent 95% confidence intervals of the mean. Promoters are putative and represent 500bp upstream of a gene without any other genomic feature overlap. (c) Upset plot showing common genes containing a hypermethylated exon per hypermethylated sex/caste per comparison. The set size indicates the total number of hypermethylated genes, the intersection size shows how many of those are common between sets, as indicated by the connections in the bottom panel. E.g. 34 genes are uniquely hypermethylated in males compared to workers.

Genome-wide, we see overall similar levels of DNA methylation across various genomic features for both sexes and castes (Fig.1b). It has recently been shown that promoter DNA methylation exists in some insect species (Lewis *et al*., 2020; Bain *et al*., 2021). We have, therefore, annotated putative promoter regions in *B. terrestris*, defined at 500bp upstream of a gene with no overlap with other genomic features, we also added UTR regions and intergenic regions to further explore the genome-wide methylation profile. We find the highest levels of DNA methylation for all sexes and castes are within exon regions, whilst promoter, and 5’ UTR regions show a depletion in DNA methylation compared to intergenic regions (Fig.1b).

We also segregated genes into categories of differing levels of DNA methylation to explore the potential function of highly methylated genes across sexes and castes. There are a small number of genes classed as highly methylated (weighted methylation level >0.7) across each sex/caste (supplementary Fig.S2, supplementary 1.0.3). Most highly methylated genes in queens and workers are also found in another caste/sex. Whereas males show a larger number of unique genes which are highly methylated (supplementary Fig.S2, n = 17). We then carried out an gene ontology enrichment test for each list of highly methylated genes per sex/caste and compared these to lists of genes classed as methylated (i.e. a weighted methylation level across the gene greater than the lambda conversion rate) for each sex/caste. We find a variety of GO terms enriched across sexes and castes mostly involved in core cellular processes (supplementary 1.0.4). Male highly methylated genes did, however, have a few GO terms enriched for eye pigmentation (GO:0008057, GO:0006726, GO:0048069) which were not present in the queen and worker enriched GO terms.

### Differential DNA methylation between sexes and castes

A differential DNA methylation analysis between sexes and castes found a total of 1,232 differentially methylated CpGs between males and reproductive workers, 1,034 differentially methylated CpGs between males and queens and 358 differentially methylated CpGs between queens and reproductive workers. Roughly equal numbers were hypermethylated in each sex/caste per comparison, except for males and queens where queens show slightly more hypermethylated sites (Chi-squared Goodness of Fit: χ^2^ = 11.28, *df* = 1, *P* < 0.01, male n = 463, queen n = 571). The majority of all differentially methylated CpGs are located within genes and specifically within exons, we also find a slight depletion of differentially methylated CpGs in the first exon compared to the following exons (supplementary Fig.S1), this is in line with DNA methylation being slightly lower in the first exon in *B. terrestris* (Lewis *et al*., 2020).

We next classed a gene as differentially methylated if a given exon contained at least two differentially methylated CpGs and had an overall weighted methylation difference of at least 15%. We find 161 genes are differentially methylated between males and workers and males and queens (with some differences between the lists) and 59 between queens and workers (supplementary 1.0.5). Previous research has suggested there are two classes of methylated genes in arthropods, highly methylated genes involved in housekeeping functions and genes with lower levels of DNA methylation which are more plastic (e.g. Asselman *et al*., 2016). We find almost all genes classed as differentially methylated in this study do indeed show low levels of DNA methylation (>0 and <0.3) in all three castes, with a few showing medium levels (>=0.3 and <0.7) and none being classed as highly methylated genes in any sex/caste (supplementary Fig.S2).

We carried out a GO enrichment analysis on all differentially methylated genes and on hypermethylated genes for each sex/caste per comparison (supplementary 1.0.6). Whilst most terms are involved in core cellular processes, we specifically find differentially methylated genes between queens and workers are enriched for chromatin-related terms (e.g. “*histone H3-K27 acetylation*” (GO:0006338) and “*chromatin remodeling*” (GO:0097549)) and reproductive terms (e.g. “*germ cell development*” (GO:0007281) and “*sexual reproduction*” (GO:0019953)).

Differentially methylated genes between males and workers were also enriched for a large number of histone modification related terms (e.g. “*regulation of histone H3-K9 methylation*” (GO:1900112), “*regulation of histone deacetylation*” (GO:0031063)) as well as reproductive related terms (e.g. “*gamete generation*” (GO:0007276), “*oviposition*” (GO:0018991) and “*spermatogenesis*” (GO:0007283)). Multiple histone related terms and reproductive terms were also found for differentially methylated genes between males and queens, as well as the above we also found “*histone H4-K20 demethylation*” (GO:0035574), “*histone H3-K27 acetylation*” (GO:0043974), “*histone H3-K27 demethylation*” (GO:0071557) and “*positive regulation of histone H3-K9 trimethylation*” (GO:1900114). When looking specifically at hypermethylated genes per sex/caste compared to all differentially methylated genes per comparison we find various regulatory terms throughout, including terms involved in hormone regulation enriched in male hypermethylated genes compared to queens (supplementary 1.0.6).

Around 66% of all differentially methylated genes occur uniquely between comparisons, with 69/205 (∼ 33%) genes occurring in multiple comparisons (Fig.1c). Specifically, we find 21 genes are hypermethylated in queens and workers when compared to males and 20 genes are hypermethylated in males when compared to queens and workers. We carried out a GO enrichment on these genes using all differentially methylated genes from all comparisons as a background set. We find general cellular processes enriched in both gene lists with hypermethylated genes in the males compared to the female castes also enriched for hormone regulatory processes (e.g. “*regulation of hormone levels*” (GO:0010817) and “*hormone secretion*” (GO:0046879)).

### Relationship between the genome and the methylome

Recent research has indicated that the underlying genome may play a role in the establishment of DNA methylation profiles in various insect species (e.g. Wang *et al*., 2016; Yagound *et al*., 2020), including *B. terrestris* (Marshall *et al*., 2019). However, the function of gene-body DNA methylation in insects remains unknown (Provataris *et al*., 2018). It has been suggested that DNA methylation induced by the environment may either be a direct substrate for selection or itself induce genetic variation (via the deamination of cytosines to thymines), creating phenotypic variation (Skinner and Nilsson, 2021).

To explore the relationship between the genome and methylome in this context, we used whole genome re-sequencing data from the males and queens to see if DNA methylation appears to be directly associated with the underlying genome. We determined the proportion of zero-fold and four-fold degenerate sites which are classed as methylated (via the binomial test described above) to see if DNA methylation is enriched at sites which may be subject to selection (zero-fold sites, which always result in an amino acid change when there is a polymorphism) compared to sites protected from selection (four-fold sites, for which any polymorphism would result in the same amino acid).

We find that a higher proportion of zero-fold degenerate sites are methylated compared to two-, three- and four-fold sites (linear model, *F*_7,24_ = 8.426, *t* = 21.862, *P* <0.001, Fig. 2a), with queens showing a higher proportion of methylated sites across all degeneracy-levels compared to males (linear model, *F*_7,24_ = 8.426, *t* = 2.337, *P* = 0.028, Fig. 2a). It’s worth noting that the zero-fold degenerate sites are only found in codon positions one and two, and so by association these positions have a higher proportion of methylated sites compared to the third codon position (supplementary 1.0.7). We also had enough alleles in our study (n = 12, four diploid queens and four haploid males) to carry out a pN/pS analysis to look for signatures of selection across the genome. We do not find any association with male or female levels of DNA methylation and potential selective pressure (linear model, *F*_1,15330_ = 1.449, *t* = −1.204, *P* = 0.228, Fig.2b).

**Figure 2:**
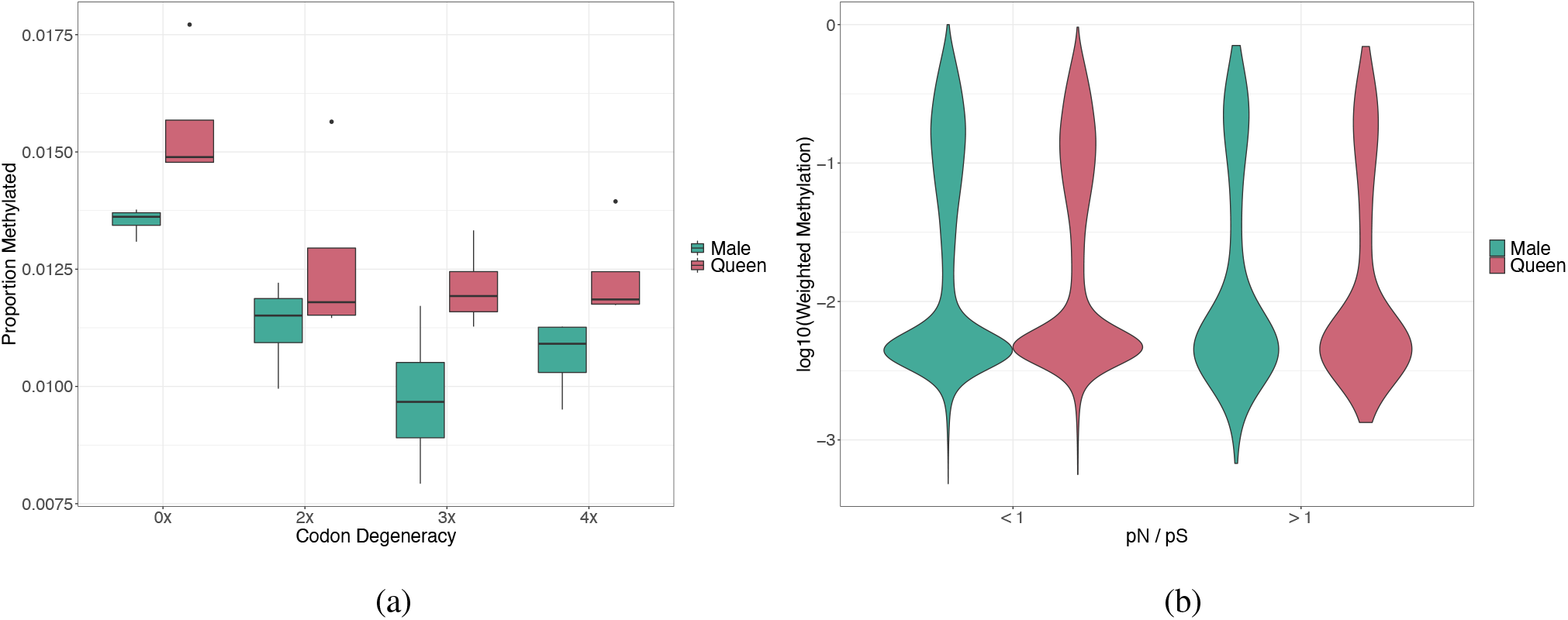
(a) Boxplot showing the proportion of methylated sites per codon degeneracy level by sex. With zero-fold degenerate sites exposed to selection and four-fold degenerate sites shielded from selection. Outliers are show by black points. (b) Violin plot, showing data via a mirrored density plot, of the DNA methylation levels, by sex, of all genes with a pN/pS >1 (i.e. genes under positive selection, n = 398) and a pN/pS <1 (i.e genes under stable selection, n = 14,934).

## Discussion

In this study we have explored the sex- and caste-specific DNA methylation profiles of *B. terrestris* and examined the relationship between the methylome and underlying genome. We find that sexes and castes show similar genome-wide DNA methylation profiles, with more variability in males. We also find there are a number of genes which are differentially methylated between sexes and castes, mostly between males and the female castes, involved in the regulation of histone modifications and reproductive-related processes. These differentially methylated genes show low levels of DNA methylation generally in all castes. Finally, we find that DNA methylation is enriched at sites exposed to selection in *B. terrestris*, i.e. zero-fold degenerate sites. However, we did not find an association between DNA methylation and genes potentially under selection.

Genome-wide we found that males show similar DNA methylation profiles to the two female castes, in terms of DNA methylation localisation to exons and depletion in promoter regions, we also found a number of differentially methylated genes between males and females. Specifically, differentially methylated genes are enriched for many histone modification related processes. It has recently been found in the silk moth that the presence of DNA methylation promotes histone H3-K27 acetylation which changes the chromatin formation of a region allowing changes in gene expression (Xu *et al*., 2021). The relationship between DNA methylation and histone modifications in social insects remains unknown. However, recent work by Choppin *et al*. (2021) shows a role for histone acetylation in the regulation of worker reproduction and gene expression in the ant *Temnothorax rugatulus*. Additionally, Dixon and Matz (2022) find gene-body methylation across a variety of invertebrates, including *B. terrestris* is not sufficient to fully explain changes in gene transcription, they also suggest a more complex regulatory mechanism may be in play involving other epigenetic modifications. An exploration of the functional relationship between DNA methylation and histone modifications is needed across a greater diversity of insect species in order to understand how these processes may interact to produce downstream gene expression and thus phenotypic differences.

We also find differentially methylated genes between queens and reproductive workers are involved in reproductive related processes. Previous work has suggested a role for DNA methylation in reproduction in *B. terrestris* (Amarasinghe *et al*., 2014), as well as other social insects (Wang *et al*., 2020; Bonasio *et al*., 2012), although this does not appear to be consistent across Hymenoptera (Libbrecht *et al*., 2016; Patalano *et al*., 2015). Whilst the differentially methylated genes identified here suggest a role for DNA methylation in maintaining or generating sex and caste differences, a direct causal link between DNA methylation and the gene expression changes mediating phenotypes has yet to be found. The development of CRISPR-Cas9 DNA methylation editing (Vojta *et al*., 2016) in *B. terrestris* would allow for studies into the causal consequences of gene methylation.

Whilst the function of gene-body DNA methylation remains unknown, higher levels have been associated with stable gene expression of housekeeping genes (e.g. Provataris *et al*., 2018). Reciprocally, previous work suggested that low methylation of genes allows for more variation in gene expression levels increasing phenotypic plasticity (Roberts and Gavery, 2012). We find evidence in support of this idea in that all of the genes found to be differentially methylated between castes and sexes show overall low levels of DNA methylation, with no highly methylated genes differentially expressed. However, it is worth noting that there are proportionally more lowly methylated genes in this species compared to highly methylated genes which may also account for this trend. In addition, whilst the differential DNA methylation analysis done here is particularly stringent for this field, requiring a percentage difference in DNA methylation levels will allow for more lowly methylated genes to be identified, compared to highly methylated genes which would require overall greater differences in DNA methylation to meet the 15% threshold. Nevertheless, there is some mixed evidence across invertebrates that suggests lowly methylated genes respond to environmental changes as a group by increasing methylation levels and thus decreasing transcriptional levels, with the opposite occurring for highly methylated genes, providing transcriptional plasticity, known as the ‘seesaw’ hypothesis (Dixon *et al*., 2018; Dixon and Matz, 2022). A re-analysis of reproductive and sterile bumblebee worker WGBS data as part of Dixon and Matz (2022), found some support for this theory. A study specifically addressing this idea, which is able to disentangle tissue-specific profiles is needed.

We next assessed the relationship between the underlying genome and the methylome. As discussed in Dixon and Matz (2022), gene-body DNA methylation does not appear to directly regulate gene expression in invertebrates. However, recent papers have suggested a role for DNA methylation in species evolution, whereby environmentally induced DNA methylation is directly heritable (Skinner and Nilsson, 2021; Harney *et al*., 2022) and may produce adaptive phenotypes. Multiple studies in arthropods have indeed demonstrated heritability of DNA methylation (e.g. Wang *et al*., 2016; Yagound *et al*., 2020) including environmentally induced DNA methylation (Harney *et al*., 2022). Given that DNA methylation has also been shown to be linked to the underlying genotype in various insect species (Wang *et al*., 2016; Marshall *et al*., 2019; Yagound *et al*., 2019, 2020; Wu *et al*., 2020) we predicted that it may be enriched in certain codon positions in order to provide substrate for selection, not through directly altering gene expression levels but through its mutagenic effect on the underlying genome, which may then cause transcriptional variation.

In order to begin to explore this idea in *B. terrestris*, we examined the relationship between DNA methylation and codon degeneracy. We found a higher proportion of zero-fold degenerate sites (those exposed to selection) are methylated compared to two-fold, three-fold and four-fold sites. This trend is also observed in humans (Chuang and Chen, 2014). Given that the presence of DNA methylation has been shown, in humans, to increase mutation rates by over ten-fold compared to background genomic rates via the spontaneous deamination of cytosines (Sved and Bird, 1990), we believe this finding provides some preliminary support for the idea of DNA methylation influencing gene expression changes via the underlying genome. However, Chuang and Chen (2014) argue that enrichment of DNA methylation at zero-fold degenerate sites is not to act as a mutagen for such sites but rather to serve as a stabilising factor for these sites. Given that we find genes with high levels of gene-body DNA methylation are involved in housekeeping functions and stable gene expression, this would support this idea. As mentioned above, however, there is now a thought that there are two classes of methylated genes, those highly methylated with housekeeping functions and those lowly methylated which may be more plastic.

Finally, we did not find any relationship between DNA methylation levels of a gene and the pN/pS ratio. This may be due to a number of reasons. Firstly, there may simply be no relationship between DNA methylation and rates of positive selection in *B. terrestris*, countering the idea that DNA methylation is providing substrate for selection. Secondly, the samples used in this study are from commercial colonies and as such may not be under any particular environmental pressures driving selection. Exploring this relationship in natural populations would provide a better indicator for a potential role of DNA methylation in adaptive processes. Additionally, Park *et al*. (2011) found the proxy for DNA methylation (CpG observed / expected density) was negatively correlated with rates of both synonymous and non-synonymous substitutions between species of the parasitic wasp *Nasonia*. Again, this supports the idea that methylation is present in constrained genes, potentially acting as a stabilising factor. However, as mentioned above this needs to be assessed fully in terms of highly methylated and lowly methylated genes rather than on a genome-wide scale in order to fully understand this relationship.

## Conclusion

We have characterised the sex- and caste-specific methylome of *B. terrestris* identifying a potential role for DNA methylation in downstream epigenetic regulatory processes which may influence sex and caste phenotypic differences. These results are correlational but can direct future experimental manipulation of DNA methylation profiles at specific genes. We also find that differentially methylated genes are those showing low overall levels, this may be due to the nature of the statistical analysis used to identify differential DNA methylation or it could suggest a function for low-levels of DNA methylation in mediating plasticity of gene expression. We also find that DNA methylation is enriched at zero-fold degenerate sites and suggest this may be explained by DNA methylation functioning as a mutagen at these sites to provide substrate for selection via changes in gene transcription mediated by the underlying genotype.

## Acknowledgements

We thank the editors and reviewers for their helpful feedback on earlier versions of this manuscript. We thank Biobest N.V. for providing us with the bumblebee colonies and Dr Ben Hunt and Dr Andrew Mongue for analysis advice and Dr Christian Thomas for shipping the bees from Leicester to Edinburgh. H.M. had a PhD-scholarship from the Central England NERC Training Alliance (NERC, UK). This study was funded in the context of the NERC grant NE/N010019/1 and a Leverhulme Trust Research Project Grant (RPG-2020-363) awarded to E.B.M., the KU Leuven BOF Centre of Excellence Financing on ‘Eco- and socio-evolutionary dynamics’ (Project number PF/2010/07) and a grant from the Research Foundation-Flanders (FWO-Vlaanderen, grant G.0463.12) awarded to T.W. and the European Research Council grant: PGErepro awarded to L.R. This research used the ALICE2 High Performance Computing Facility at the University of Leicester.

## Author contributions

E.B.M., T.W. and H.M. conceived the study. Colonies were generated by Biobest (Westerlo, Belgium) under supervision of F.W. J.S.Z. carried out the dissections to confirm the reproductive status of workers. H.M. carried out the wet lab work. H.M. and M.T.N carried out the analyses. L.R. contributed the whole genome re-sequencing. H.M. wrote the initial manuscript. All authors contributed to and reviewed the final manuscript.

## Data Accessibility

Data has been deposited in GenBank under NCBI BioProject: PRJNA779586. All code is available at: https://github.com/MooHoll/Parent_of_Origin_Methylation.

