## Supplementary 2 for "Caste- and sex-specific DNA methylation in a bumblebee is associated with codon degeneracy"

### Supplementary figures and text

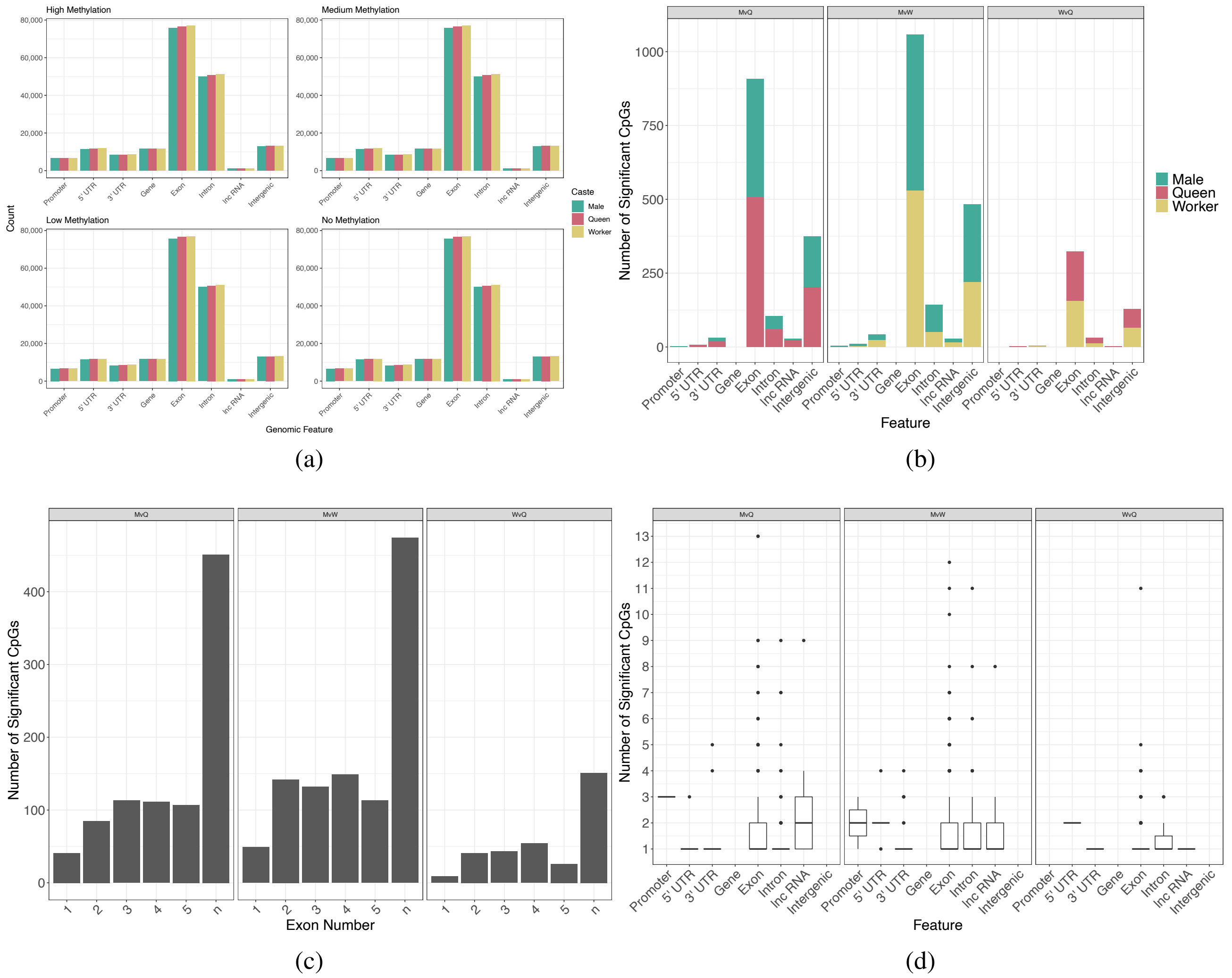


Figure S1: (a) Bar plots of the total number of genomic features categorised by weighted methylation level for sexes and castes. High methylation is a weighted methylation level >0.7, medium is >0.3-0.7, low is >0-0.3 and no methylation is equal to zero. (b) Staked bar chart showing the genomic location of the differentially methylated CpGs sites per comparison, coloured by the hypermethylated caste. (c) Barplot of the number of significant CpGs in the first five exons, with 'n' representing exons six on-wards. (d) Boxplots of the number of significant CpGs per feature, per comparison, each dot represents an outlier.

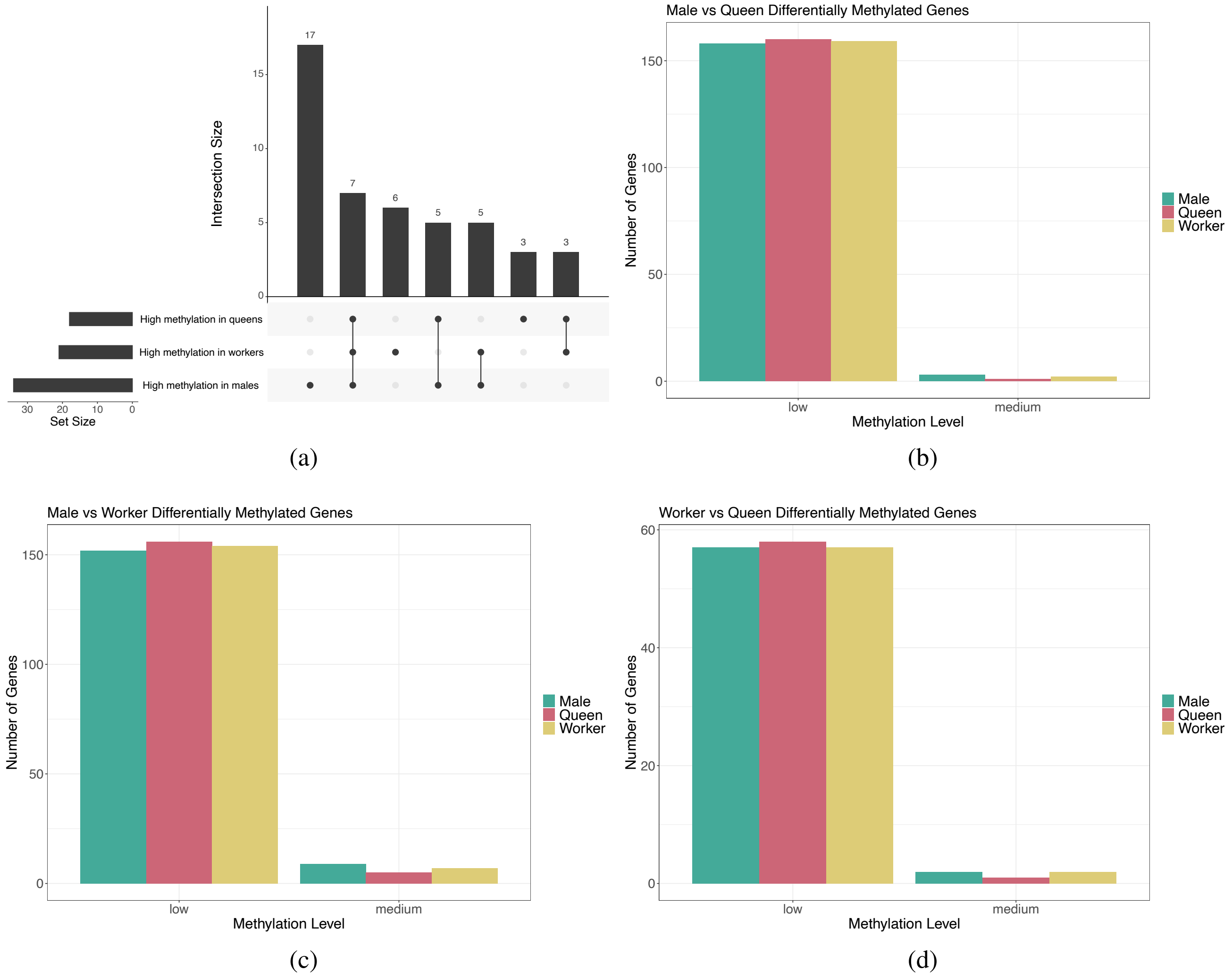


Figure S2: (a) UpSet plot showing the common genes which are highly methylated (>0.7 weighted methylation level) per sex/caste. The set size indicates the number of highly methylated genes, the intersection size shows how many of those are common between sets or unique, as indicated by connections in the bottom panel. (b - d) Overall methylation level of genes per caste which are differentially expressed between castes, for male vs queens, male vs workers and workers vs queens respectively. Low methylation refers to an overall weighted methylation >0 and <0.3, medium methylation refers to >= 0.3 and <0.7.
